# Mapping binding landscape of allosteric inhibitor G6PDi-1 on human G6PD

**DOI:** 10.64898/2026.04.21.719807

**Authors:** Amit Kumawat, Andrea Perra, Marina Serra, Giorgia Zedda, Marta Anna Kowalik, Andrea Marco Caddeo, Paolo Ruggerone

**Author notes:** **Corresponding Author:** Amit Kumawat.

## Abstract

Targeting the oxidative pentose phosphate pathway by inhibiting glucose-6-phosphate dehydrogenase (G6PD), is a promising anticancer strategy, yet clinically useful inhibitors remain unavailable. A key limitation is the lack of molecular insight into how allosteric and non-competitive inhibitors perturb enzymatic activity, limiting rational optimization. Here, we investigated the mechanism of G6PDi-1, a potent, reversible, non-competitive G6PD inhibitor, by combining biochemical assays with molecular dynamics simulations, Markov state model and MM/PBSA binding energy calculations. Experimentally, G6PDi-1 reduced active dimer formation in hepatoblastoma HepG2 cells and increased the inactive monomer fraction, linking inhibition to disrupted oligomerization. Computationally, we identified multiple sites on the enzyme surface, with preferential binding at dimer interface that sterically blocks oligomerization. Importantly, additional distal sites showed enhanced inter-residue correlations, suggesting secondary allosteric effects that shift G6PD toward monomeric states less competent for oligomerization. These findings provide a molecular basis for structure-based development of improved strategies for G6PD inhibition suited for cancer therapy.

---

Glucose-6-phosphate dehydrogenase (G6PD, EC 1.1.1.49) is the rate-limiting enzyme of the oxidative pentose phosphate pathway (PPP), which generates ribose-5-phosphate for nucleotide synthesis and NADPH for reductive biosynthesis and antioxidant defense.^1,2^ G6PD expression and activity are upregulated in numerous tumors (e.g. hepatocellular carcinoma (HCC)), contributing to several hallmarks of cancer, including cell proliferation, metastasis, deregulated cellular metabolism and decreased overall survival.^3–9^ Given the key role of the PPP in cancer metabolic reprogramming, it is not surprising that targeting the pathway with specific G6PD inhibitors represents a promising therapeutic option for several cancers, including HCC.^10,11^ Despite this, no G6PD inhibitors have reached clinical use and developing drug-quality inhibitors remains burdensome and challenging. Structurally, human G6PD enzyme exists in an active form as dimer or tetramer of identical subunits, whereas the monomeric form is catalytically inactive.^12^ Each G6PD monomer contains a substrate binding site for glucose-6-phosphate and two NADP⁺ binding sites, one for catalysis and a second structural site that helps stabilize the enzyme’s oligomeric state.^13,14^

Earlier studies identified steroidal and non-steroidal small molecules (e.g. dehydroepiandrosterone (DHEA), thienopyrimidines, and benzothiazinones) as putative G6PD inhibitors in several *in vitro* and *in vivo* cancer models, but their clinical translation has been limited by the lack of target specificity and effects that may have resulted from off-target mechanisms rather than direct G6PD inhibition.^15–19^ Rabinowitz and coworkers made a significant breakthrough with the identification of the reversible and noncompetitive inhibitor G6PDi-1 (IC_50_ ≈ 0.07 μM).^20,21^ Although G6PDi-1 is suggested to act allosterically, the molecular mechanism of this effect remains unknown. It has been reported that G6PDi-1 disrupts G6PD oligomerization in human immortalized embryonic kidney HEK293T cells.^22^ We tested G6PDi-1 in human hepatoblastoma HepG2 cells (Cellosaurus RRID: CVCL_0027, DepMap ID: ACH-000739), a cell line widely used for drug safety and toxicity assays, as well as in HCC-related studies, as it displays the key features of a liver neoplastic transformation (see Supplementary Information (SI) for experimental methods).^23,24^ As evident from **Figure S1**, we observed significant reduction in the dimer formation following exposure to G6PDi-1. This finding extends the oligomer-disrupting effect of G6PDi-1 to a more disease-relevant hepatic cell context and underscores the need to elucidate the molecular basis of this inhibition to support analogue optimization and future inhibitor design.

To investigate the mechanism of G6PDi-1 mediated inhibition, we first used the Schrodinger’s SiteMap tool to identify potential cavities on G6PD based on geometric and physicochemical criteria (**Figure S2A, Table S1)**.^25^ G6PDi-1 was docked into the six identified cavities using Glide tool (**Figure S2B**),^26^ followed by rescoring of the top poses using MM-GBSA binding free energy calculations (**Table S2**).^27^ We then performed all-atom molecular dynamics (MD) simulations of the six docked complexes, each representing a distinct binding site to characterize the protein-ligand interaction landscape. For each complex, we ran four independent replicates of approximately 5 μs each, generating a total simulation time of ∼120 μs (see SI for computational methods). Our simulations revealed that G6PDi-1 interacts in a highly dynamic manner exhibiting ligand unbinding events within tens to hundreds of nanoseconds and potential reassociation with the same or alternative regions on the protein surface. We quantified these transient interactions by calculating residue-wise number of contacts between the protein and ligand (**Figure 1A**). Minimal interactions with the substrate binding site residues corroborate the experimental activity of G6PDi-1 as a non-competitive inhibitor.^20^ Seven distinct cavities (C1-C7) were defined based on residues exhibiting the highest contact frequency with G6PDi-1, representing localized clusters of frequently contacted surface regions (**Figure 1B**). C1 and C2 correspond to the region near the catalytic (NADP⁺-C) and structural (NADP⁺-S) cofactor binding sites, respectively; C3 represents the distal pocket located opposite the dimer interface; C4 marks the dimer interface binding region; C5, C6 and C7 denote intermittent sites identified from residues with the highest contact counts. However, the frequent transitions between these sites prevented the identification of a single dominant binding pocket based solely on contact analysis or visual inspection of the trajectories.

**Figure 1.**
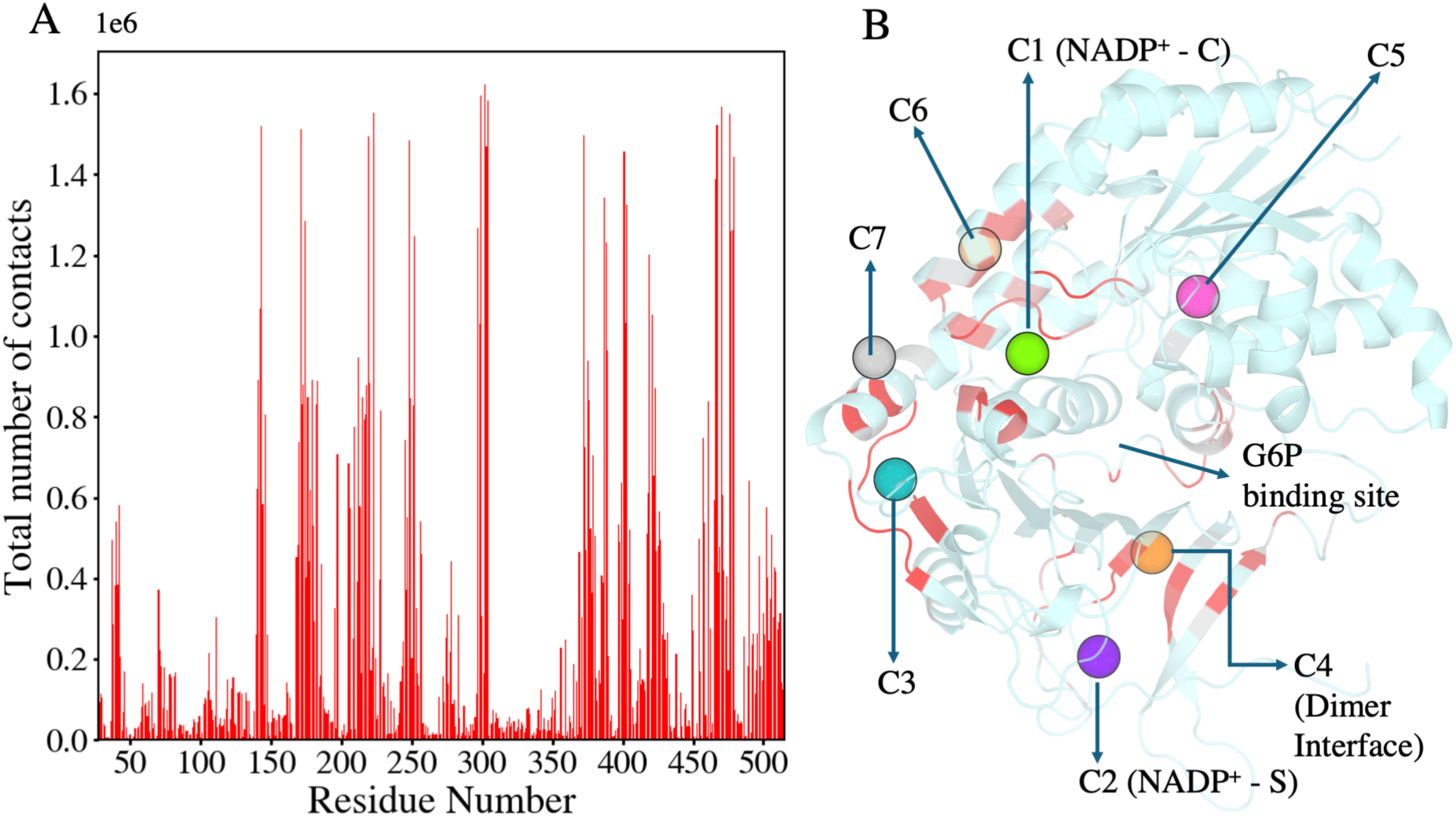
Residue-wise contact analysis and definition of G6PDi-1 binding sites on G6PD. (A) Residue-wise contact plot shows the total number of contacts between G6PDi-1 and G6PD residues observed during the simulations. A contact was defined when the distance between the center of mass (COM) of the ligand and the COM of a residue was less than 0.45 nm. (B) Structural mapping of ligand binding sites on G6PD from the contact analysis. The protein is shown in cartoon representation, with residues exhibiting high contact frequency (>5 × 10^5^ frames) highlighted in red. Seven binding cavities from C1 to C7 were defined based on clusters of residues with high contact frequency.

Subsequently, these extensive transitions between cavities motivated a more quantitative kinetic description using a hidden Markov state model (HMSM) constructed from the MD trajectories (see SI for more details) using PyEMMA package.^28–32^ For this purpose, the minimum distance between the COMs of G6PDi-1 and the residues defining each cavity was used as the feature set, allowing an explicit state decomposition of the ligand interaction sites on the G6PD surface. Implied timescale analysis together with Chapman-Kolmogorov validation identified a lag time of 30 ns, and this value was used for all subsequent HMSM calculations (**Figure S3A**).^33^ The resulting kinetic network showed that unbinding from the cavities occurred on relatively similar timescales, with mean first passage times (MFPTs) for escape of approximately 20 μs across multiple sites (**Figure 2A**). In contrast, association kinetics showed site dependence, with the dimer interface cavity C4 exhibiting the fastest association, with an MFPT of ∼ 118 μs, whereas binding to the structural NADP⁺ site occurred on much slower timescales (∼ 3 ms), which is likely due to the high flexibility of the C-terminal region shaping this pocket. The ligand bound conformational ensembles therefore represent metastable states sampled on the simulation timescale. Hence, the derived association and dissociation kinetics are interpreted here primarily to identify and compare kinetically accessible binding regions, rather than as a direct quantitative estimate of experimental potency.^34^ An MFPT cutoff of 250 μs was chosen to capture the faster associating sites while excluding slower binding modes in the 0.7-3 ms range. The cavities C1 (NADP⁺ catalytic site), C3 (distal pocket), and C4 (dimer interface) were identified as kinetically favored binding regions and selected for further analysis.

**Figure 2.**
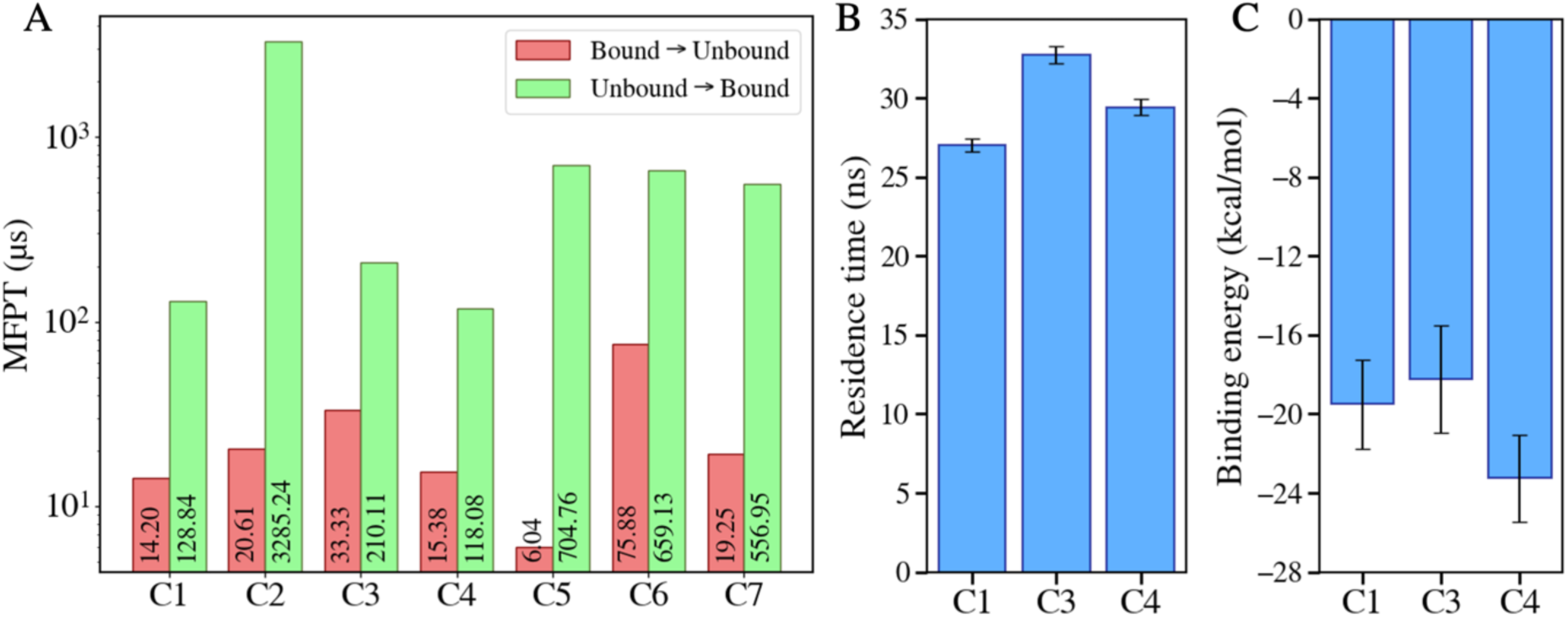
Kinetics and energetics of G6PDi-1 binding to G6PD cavities. (A) MFPTs for ligand transitions between bound and unbound states at each cavity (C1-C7), obtained from the HMSM. Red bars show MFPTs for unbinding (bound to unbound), and green bars show MFPTs for binding (unbound to bound). The logarithmic scale highlights site dependent differences in both association and dissociation kinetics, with fastest binding at the dimer-interface cavity C4 (∼ 118 µs) and slowest association at the structural NADP⁺ site C2 (∼ 3.3 ms). (B) Average ligand residence times at the three kinetically favored cavities (C1, C3, C4) derived from MD trajectories, with error bars indicating the standard error of the mean. (C) Binding free energies of G6PDi-1 at cavities C1, C3, and C4 calculated using the MM-PBSA approach on clustered bound ensembles. Error bars represent standard deviations across the sampled frames, showing the most favorable interaction energy at the dimer-interface site C4.

First, the average residence time of G6PDi-1 in these three cavities (C1, C3, C4) was determined over all trajectories. A ligand was considered bound when the distance between the COMs of G6PDi-1 and the corresponding cavity residues is less than 0.5 nm. We observed multiple binding and unbinding events and residence time distributions were fitted with a single exponential function to obtain the average residence time (τ) for each site (**Figure 2B**). G6PDi-1 remains bound for ∼27 ns and ∼29 ns at the catalytic NADP⁺ site (C1) and the dimer interface (C4), respectively, whereas the distal pocket (C3) exhibited a slightly extended occupancy of ∼33 ns. Ligand bound frames corresponding to these sites were then extracted and clustered to analyze the ligand conformations at each pocket. Representative structures from the most populated clusters showed that G6PDi-1 adopts heterogeneous conformations at C1 and C4, consistent with local protein flexibility and the solvent exposed nature of these regions (**Figure S4**). In contrast, binding at C3 was associated with a narrower distribution of ligand poses, indicative of a more constricted local environment.

Binding free energy calculations were subsequently performed using MM-PBSA approach for the frames belonging to the top clusters that cumulatively accounted for 75% of the bound ensemble at each site.^35,36^ Despite the higher ligand fluctuations and shorter residence time, the dimer interface pocket (C4) has more favorable binding energetics for G6PDi-1, followed by C1, whereas the distal pocket (C3) shows comparatively weakest interaction energy among the three sites (**Figure 2C**). Furthermore, decomposition of the binding free energies into residue-wise contributions showed that hydrophobic contacts and intermittent hydrogen bonds (H-bonds) stabilize the ligand at all three sites. (**Figure S5**). For example, at C1, both polar and nonpolar interactions contribute to ligand stabilization, including H-bonds with Tyr249 and Lys171 (**Figure S5B**). The distal pocket (C3) is characterized by a predominantly hydrophobic enclosure formed by aliphatic residues (e.g., V303, Leu468, Pro477, I480) that restrict ligand motion, together with consistent H-bond with Leu305 backbone (**Figure S5D**). Similarly, at the highest affinity dimer interface site (C4) in **Figure S5F**, binding is driven by hydrophobic interactions (e.g. Ile220, Phe373, Leu390, Met404, Leu420) combined with H-bonds involving residues such as Asn218, Asn388, Asp389, Thr402 and Ser418, which together result in the most favorable overall binding energy. This finding supports the experimental evidence of disrupted oligomerization, providing a rationale for the observed inhibition.

In order to investigate the plausible allosteric modulation through ligand binding at these sites, we compared the conformational dynamics of the ligand bound states (C1, C3, and C4) with the MD simulations trajectories (three replicates, 100 ns each) for the active tetrameric state containing substrate G6P and NADP+ at the catalytic and structural sites. **Figure 3A** shows increase in root mean square fluctuations (RMSF) in the G6PDi-1 bound systems within residues forming inter subunit contacts and dimer interface due to the disrupted inter-subunit packing as compared to the active form. We therefore next asked whether ligand binding also allosterically alters internal communication,^37–39^ and to address this we computed residue-wise normalized linear mutual information (nLMI) for the active and ligand bound systems (**Figure S6)**.

**Figure 3.**
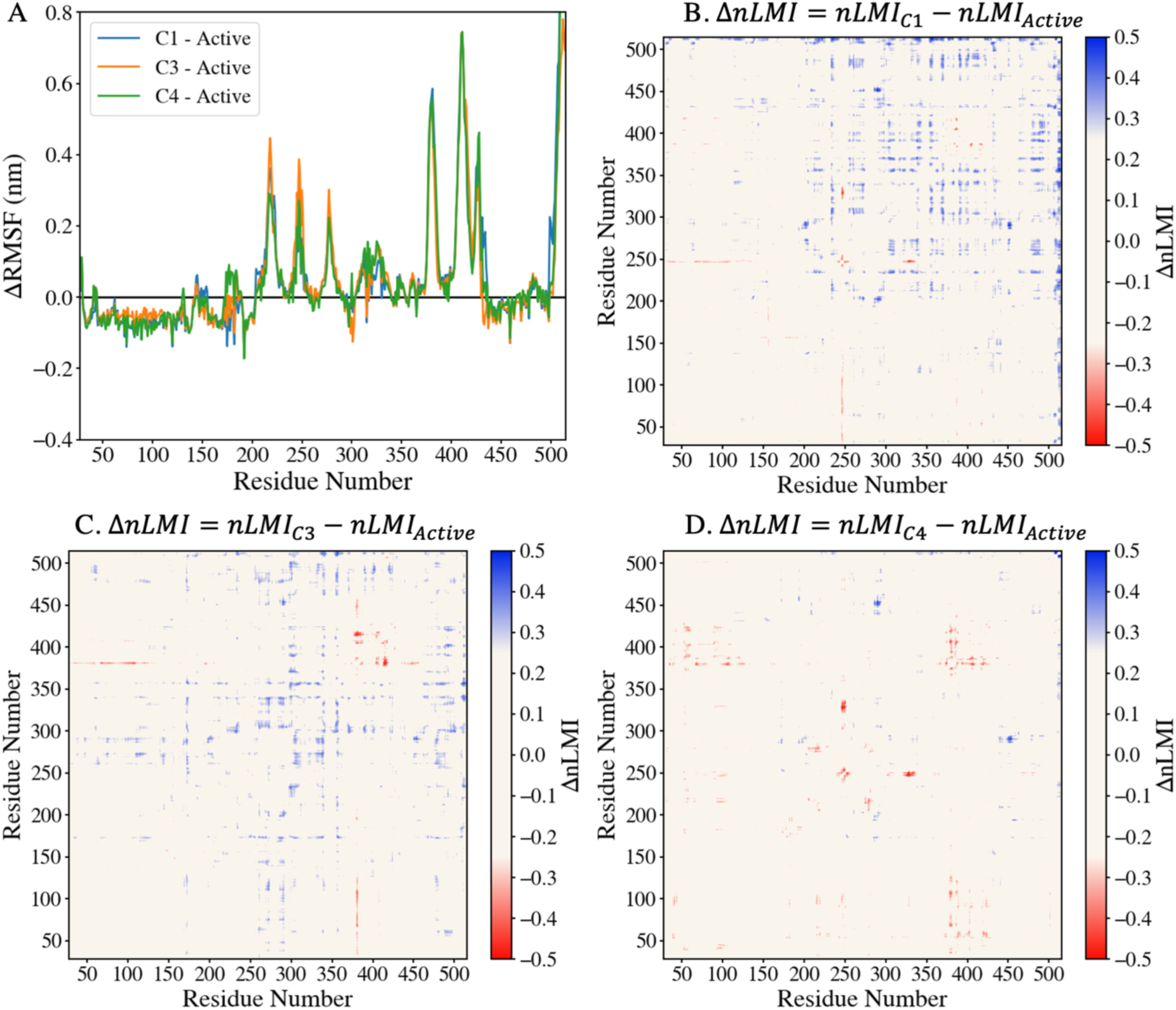
Ligand induced changes in conformational fluctuations and residue-residue coupling in G6PD. (A) Difference in RMSF between the ligand bound systems (C1, C3, C4) and the active tetrameric state. Changes were calculated as Δ*RMSF* = *RMSF*_*cavity*_ − *RMSF*_*Active*_ where the RMSF of a monomer in the tetrameric state was averaged over all subunits. (B-D) Differential residue-wise Δ*nLMI* maps comparing ligand bound states with the active form. Ligand binding at cavities C1 (B) and C3 (C) indicate enhanced inter-residue correlations relative to the active state. In contrast, the C4-bound state (D) shows only localized changes with a similar profile to the active state. Together, these results distinguish distal sites (C1, C3), which alter internal communication, from the interface-proximal C4 site, which perturbs inter-subunit packing more directly.

Figure 3B-D show the difference in the nLMI for these systems, calculated as Δ*nLMI* = *nLMI*_*cavity*_ − *nLMI*_*Active*_, where positive values (blue) indicate stronger inter-residue coupling in the ligand bound state and negative values (red) indicate stronger coupling in the active state. The nLMI matrices for ligand bound at cavities C1 and C3 show broader off-diagonal coupling than the active state (**Figure S6B, C**), and their differential maps are dominated by positive values (**Figure 3B, C**), indicating increased inter-residue coupling upon ligand binding. In contrast, the ligand bound C4 state shows a near neutral profile with only limited and localized changes (**Figure S6A, S6D and 3D**), consistent with a direct mechanism linked to the interface occlusion. Together, these results support a distinction between the sites, where ligand binding at C4 site directly interferes with oligomerization, whereas ligand binding at distal sites C1 and C3 are associated with enhanced inter-residue correlations that shift the conformational ensemble away from dimer-competent states.^40,41^

In summary, we combined biochemical data with MD simulations and binding free energy calculations to map the binding landscape of the allosteric G6PD inhibitor G6PDi-1 and define its likely mechanism of action. We demonstrate that G6PDi-1 disrupts the active dimeric form in HepG2 cells and identify the dimer interface as the most favorable binding region, consistent with a direct disruption of oligomerization. Interestingly, G6PDi-1 binding at distal sites reshape inter–residue coupling network, uncovering an additional mode of allosteric modulation. Together, these findings unearth the structural and mechanistic basis of G6PDi-1 inhibition and establish a foundation for structure-based development of new G6PD inhibitors for effective anti-tumoral therapy.

## ASSOCIATED CONTENT

The Supporting Information (Supporting_information.pdf) is available free of charge on the ACS website. The document contains experimental procedures, computational methods, Supplementary Tables S1–S2, and Supplementary Figures S1–S8.

## AUTHOR INFORMATION

### Author Contributions

A.K. performed the molecular dynamics simulations. A.K. and P.R. analyzed the computational results. M.S., G.Z., M.A.K. and A.M.C. performed and analyzed the *in vitro* experiments. A.K., A.P., and P.R., conceived and designed the study and methodology. A.P. and P.R. contributed to the funding acquisition. A.K., M.S., and M.A.K. wrote the manuscript, with contributions from all authors. All authors approved the final version of the manuscript.

### Notes

The authors declare no conflicts of interest.

### DATA AVAILABILITY STATEMENT

The MD simulation datasets that support the conclusions of this article are available in the Zenodo repository with the following doi: 10.5281/zenodo.19068230

## ACKNOWLEDGMENT

The research leading to these results was funded by: Fondazione AIRC per la Ricerca sul Cancro ETS, Grant IG 2023 ID 29155 to A.P.; European Union—NextGenerationEU through the Italian Ministry of University and Research under PNRR–M4C2-I1.3 Project PE_00000019 “HEAL ITALIA” to A.P. and P.R. CUPF53C22000750006, University of Cagliari. A.K has been funded by e.INS- Ecosystem of Innovation for Next Generation Sardinia (cod. ECS 00000038) funded by the Italian Ministry for Research and Education (MUR) under the National Recovery and Resilience Plan (NRRP) - MISSION 4 COMPONENT 2, CUP F53C22000430001 University of Cagliari, Italy. A.K. acknowledges CINECA ISCRA for access to the LEONARDO supercomputer, owned by the EuroHPC Joint Undertaking and hosted by CINECA (Italy), under the projects Class C: HP10CWQCG8 and Class B: HP10BEOUG4. The views and opinions expressed are those of the authors only and do not necessarily reflect those of the European Union, the European Commission or the Italian Ministry of the University and Research (MUR). The funders had no role in study design, data collection and analysis, decision to publish, or preparation of the manuscript.

## Materials and methods

1. **Experimental details:**

a. Cell cultures and G6PDi-1 treatment
b. Protein extraction and sample preparation
c. Western blotting analysis
2. **2. Computational details:**

a. Binding site prediction and molecular docking protocol
b. G6PDi-1 molecule parameterization
c. Molecular dynamics simulations of ligand bound G6PD
d. Molecular dynamics simulations of wild type tetrameric G6PD
e. MSM estimation and validation

## Supplementary tables (S1 and S2) with legends

**Table S1.** Scoring of G6PD binding pockets identified by SiteMap analysis using Maestro.

| Title | SiteScore | Exposure | Philic | Phobic | Volume (Å<sup>3</sup>) | Residue Number |
| --- | --- | --- | --- | --- | --- | --- |
| Site 1 | 0.97 | 0.74 | 0.40 | 0.60 | 206.49 | 58,60,206,209,210,212,213,216,217,275,276,278,279,424,432,434,438,439,442,443,448,450,451 |
| Site 2 | 0.97 | 0.59 | 0.75 | 0.63 | 177.12 | 175,251,252,253,254,256,257,329,332,333,334,335,469,472,473,476 |
| Site 3 | 0.97 | 0.74 | 0.35 | 0.57 | 295.45 | 206,210,213,214,218,220,221,224,228,229,373,374,375,376,377,381,388,400,401,402,404,405,406,407,415,420,422,424,428,431,433 |
| Site 4 | 0.96 | 0.62 | 0.65 | 0.33 | 833.02 | 38,40,41,42,43,46,47,54,57,72,73,87,88,89,112,141,142,143,144,145,146,148,170,171,172,174,175,179,182,183,201,202,205,206,237,239,241,243,244,246,249,250,252,253,258,259,263,360,365,395,396,397,398,425,431,432,433,434,435,436,437 |
| Site 5 | 0.95 | 0.76 | 0.39 | 0.51 | 118.55 | 300,301,303,304,305,468,471,472,475,477,478,479,480,494,495,498 |
| Site 6 | 0.93 | 0.67 | 0.68 | 0.25 | 627.26 | 236,238,357,363,364,365,366,368,370,372,386,387,389,393,396,397,401,403,406,409,416,417,418,419,421,422,423,426,427,429,487,493,494,496,497,501,502,503,504,505,506,507,508,509,510,511,512,513,515 |

**Table S2.**
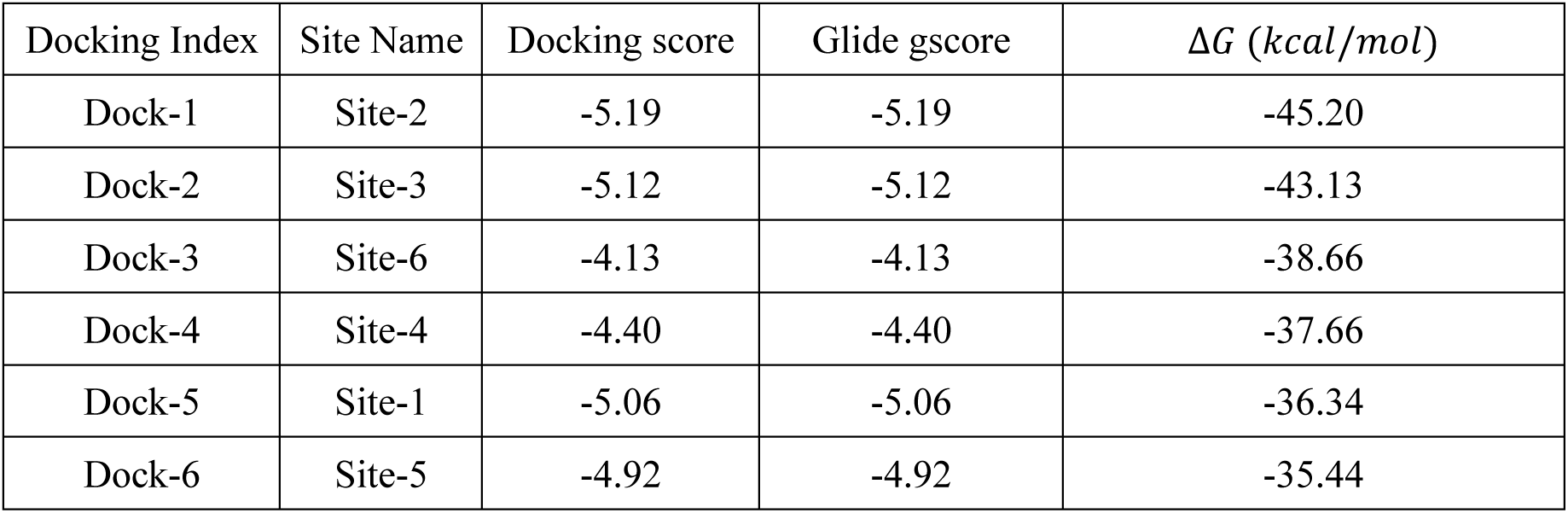
Docking scores and MM-GBSA scoring for the best docked pose of G6PDi-1 in each SiteMap identified pocket. The table is organized with docking poses ranked by most favorable MM-GBSA binding energy (*kcal/mol*).

| Docking Index | Site Name | Docking score | Glide gscore | $\Delta G$ (*kcal/mol*) |
| --- | --- | --- | --- | --- |
| Dock-1 | Site-2 | -5.19 | -5.19 | -45.20 |
| Dock-2 | Site-3 | -5.12 | -5.12 | -43.13 |
| Dock-3 | Site-6 | -4.13 | -4.13 | -38.66 |
| Dock-4 | Site-4 | -4.40 | -4.40 | -37.66 |
| Dock-5 | Site-1 | -5.06 | -5.06 | -36.34 |
| Dock-6 | Site-5 | -4.92 | -4.92 | -35.44 |

## Supplementary figures (S1 to S8) with legends

**Figure S1.**
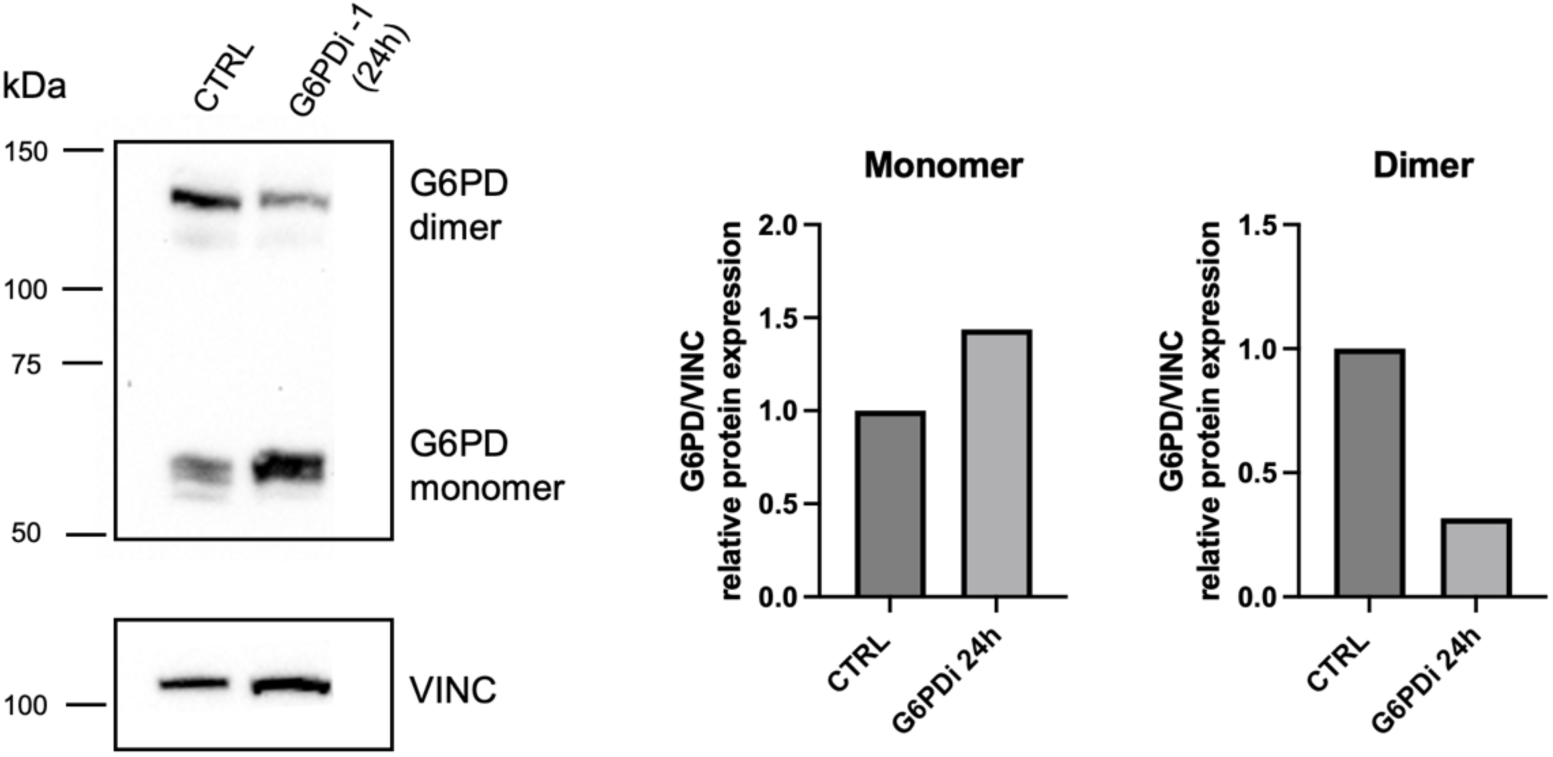
G6PDi-1 blocks formation of G6PD dimer in hepatoblastoma HepG2 cells. HepG2 cells were treated with G6PDi-1 at the concentration of 100 µM for 24 hours. Control cells (CTRL) were treated with DMSO. Western Blot of G6PD protein levels. Vinculin (VINC) was used as loading control. Western blot quantification was calculated using ImageJ software.

**Figure S2.**
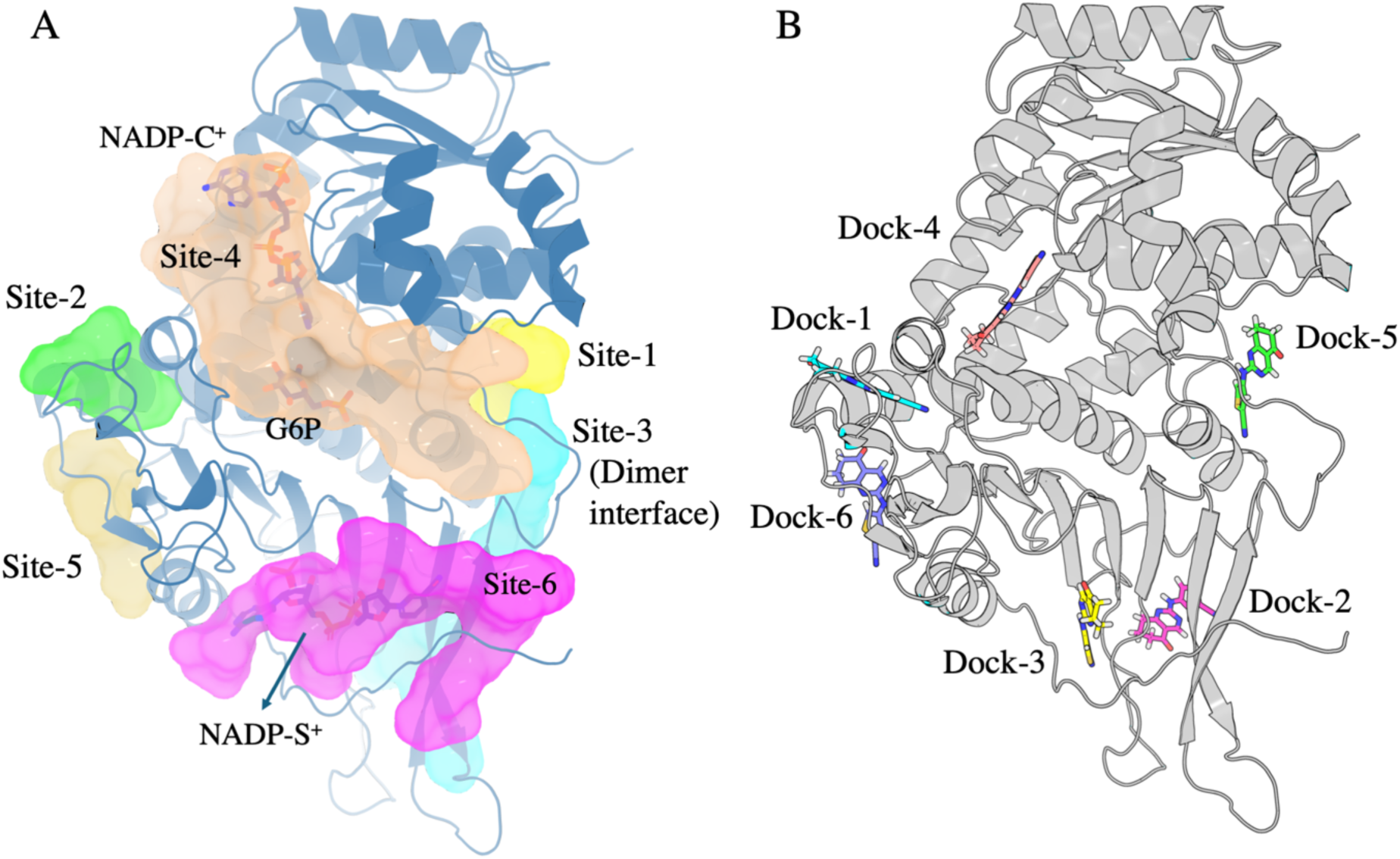
Identified binding pockets and top docked poses of G6PDi-1 on G6PD. (A) Binding pockets identified by SiteMap (Site-1 to Site-6) mapped onto the G6PD monomer crystal structure. The protein is shown in cartoon representation, with SiteMap cavities rendered as coloured surfaces. The catalytic NADP⁺ (NADP-C), structural NADP⁺ (NADP-S), and the glucose-6-phosphate (G6P) substrate are labelled for reference. The dimer interface region is highlighted. Site-5 corresponds to a distal pocket located on the opposite side of the protein relative to the dimer interface. (B) Top docked poses of G6PDi-1 (stick representation) obtained from docking against the SiteMap identified cavities. Docked poses (Dock-1 to Dock-6) are ranked according to their MM-GBSA binding energies. The association between docked poses and binding sites is reported in Table S2. These docked complexes were used as starting structures for subsequent MD simulations.

**Figure S3.**
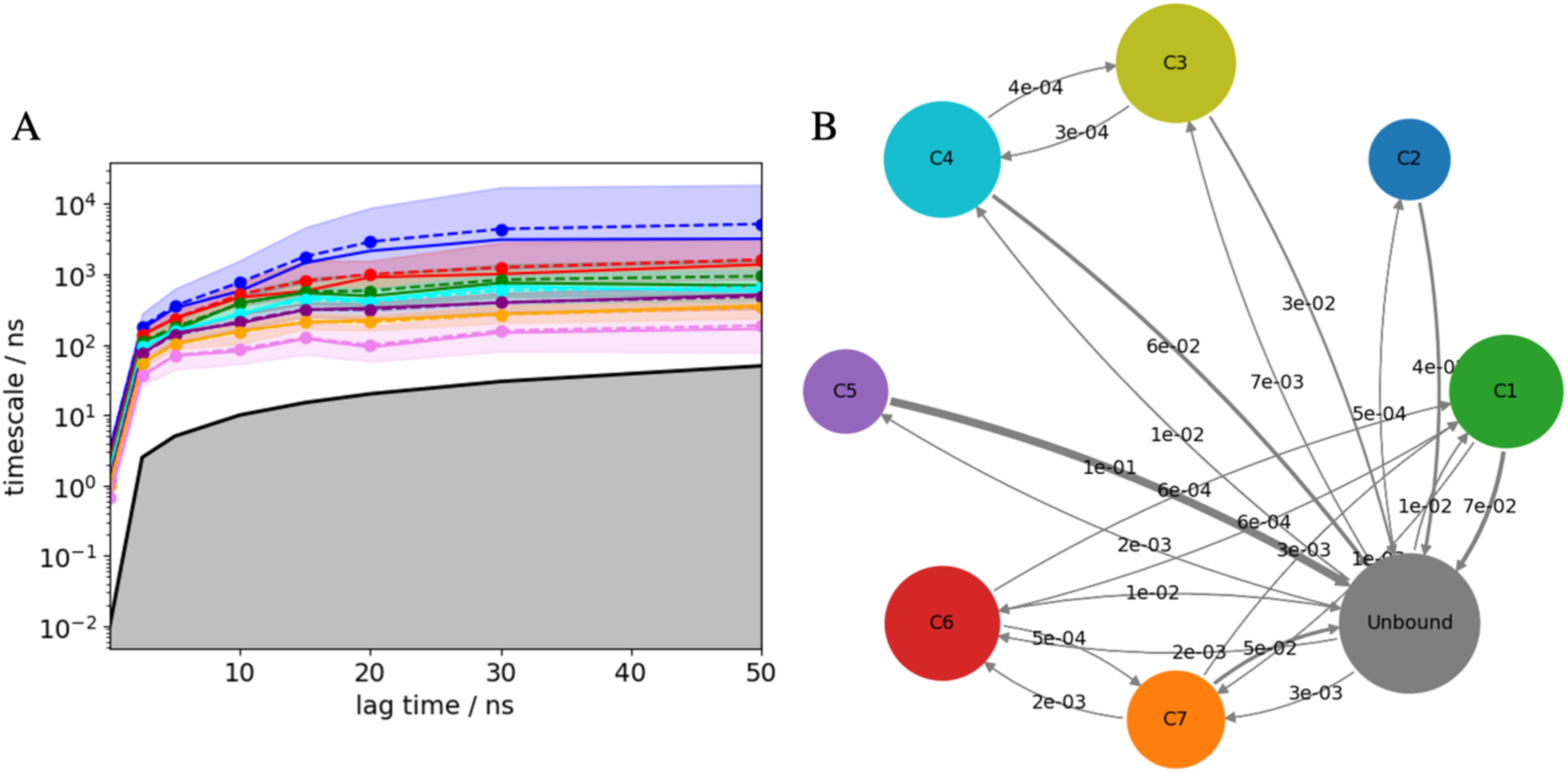
Hidden Markov state model (HMSM) of G6PDi-1 cavity exchange on G6PD. (A) Implied timescales as a function of lag time for the eight-state model support the choice of 30 ns lag time; shaded regions denote uncertainties. (B) Kinetic network connecting the seven cavity-bound states (C1–C7) and the unbound state, where node size represents stationary population and edge thickness reflects transition probability at the chosen lag time, highlighting the unbound ensemble as the central hub for ligand exchange.

**Figure S4.**
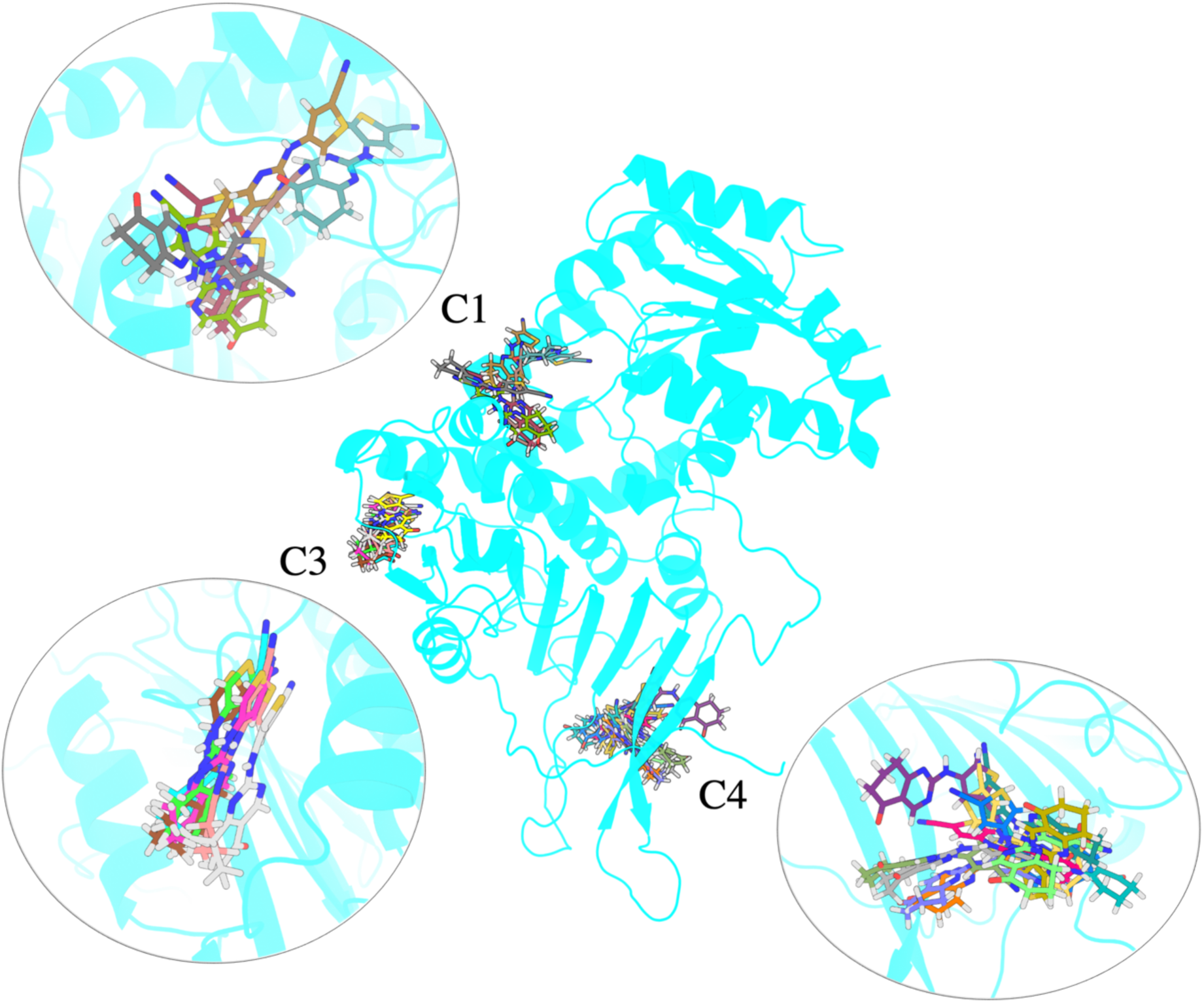
Representative conformations of G6PDi-1 at the binding cavities. G6PD monomer is shown in cartoon and representative ligand poses from the most populated clusters are shown as sticks at the C1, C3, and C4 sites. Clustering was performed on the protein backbone using ligand bound frames from the MD trajectories. Insets show enlarged images of each binding region. The ligand samples multiple conformations at C1 and C4, consistent with the more solvent-exposed and flexible pockets, whereas C3 shows a rigid distribution of poses, indicating a more confined cavity.

**Figure S5.**
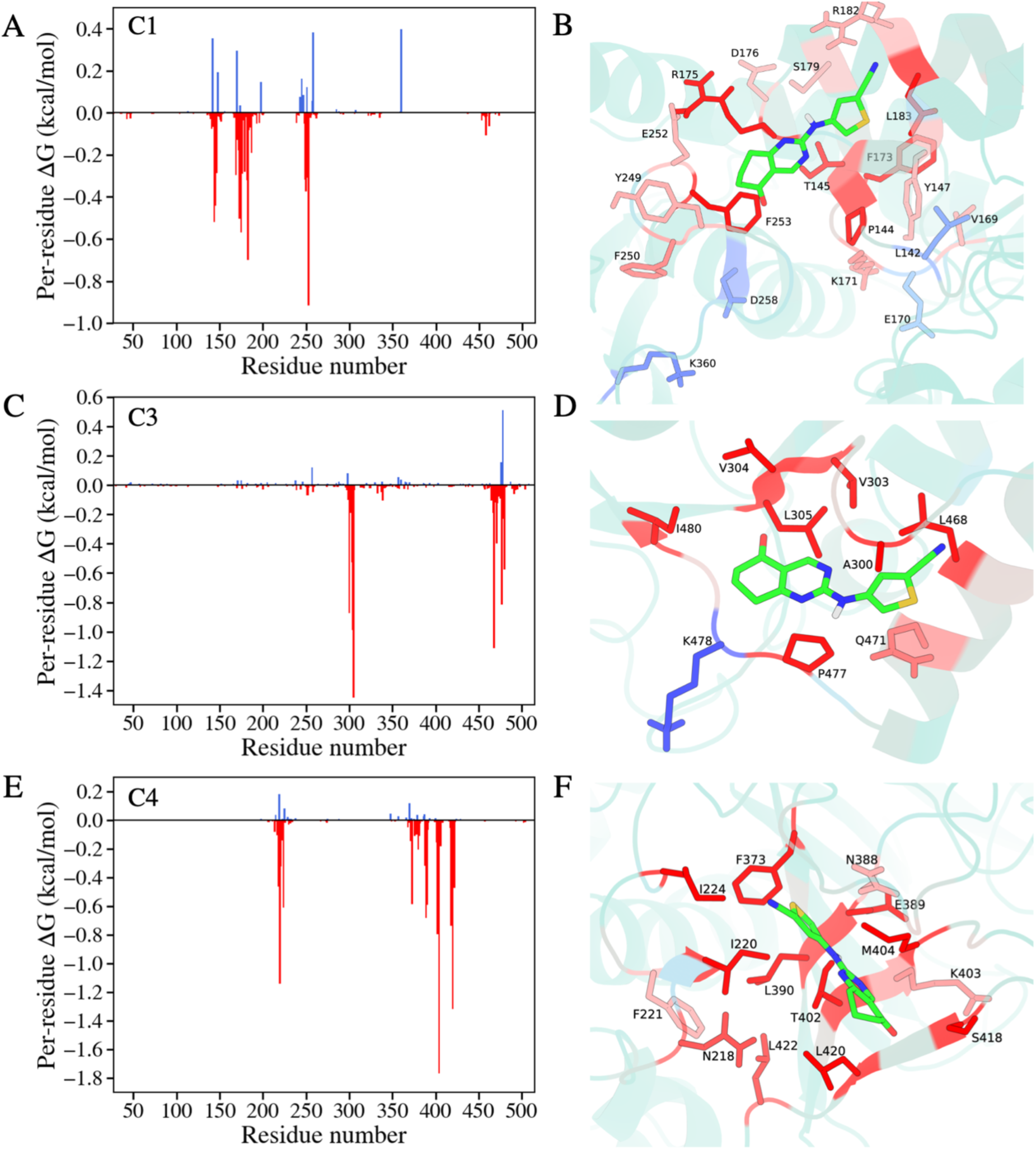
Residue-wise decomposition of the MM/PBSA binding free energy for G6PDi-1 bound at cavities C1, C3, and C4. The plots on the left show per-residue contributions for ligand bound at C1, C3, and C4, with favorable (negative) interactions shown in red and unfavorable (positive) interactions in blue. On the right, we show representative binding poses of G6PDi-1 at each cavity together with the surrounding residues, using the same color scheme as in the corresponding plots.

**Figure S6.**
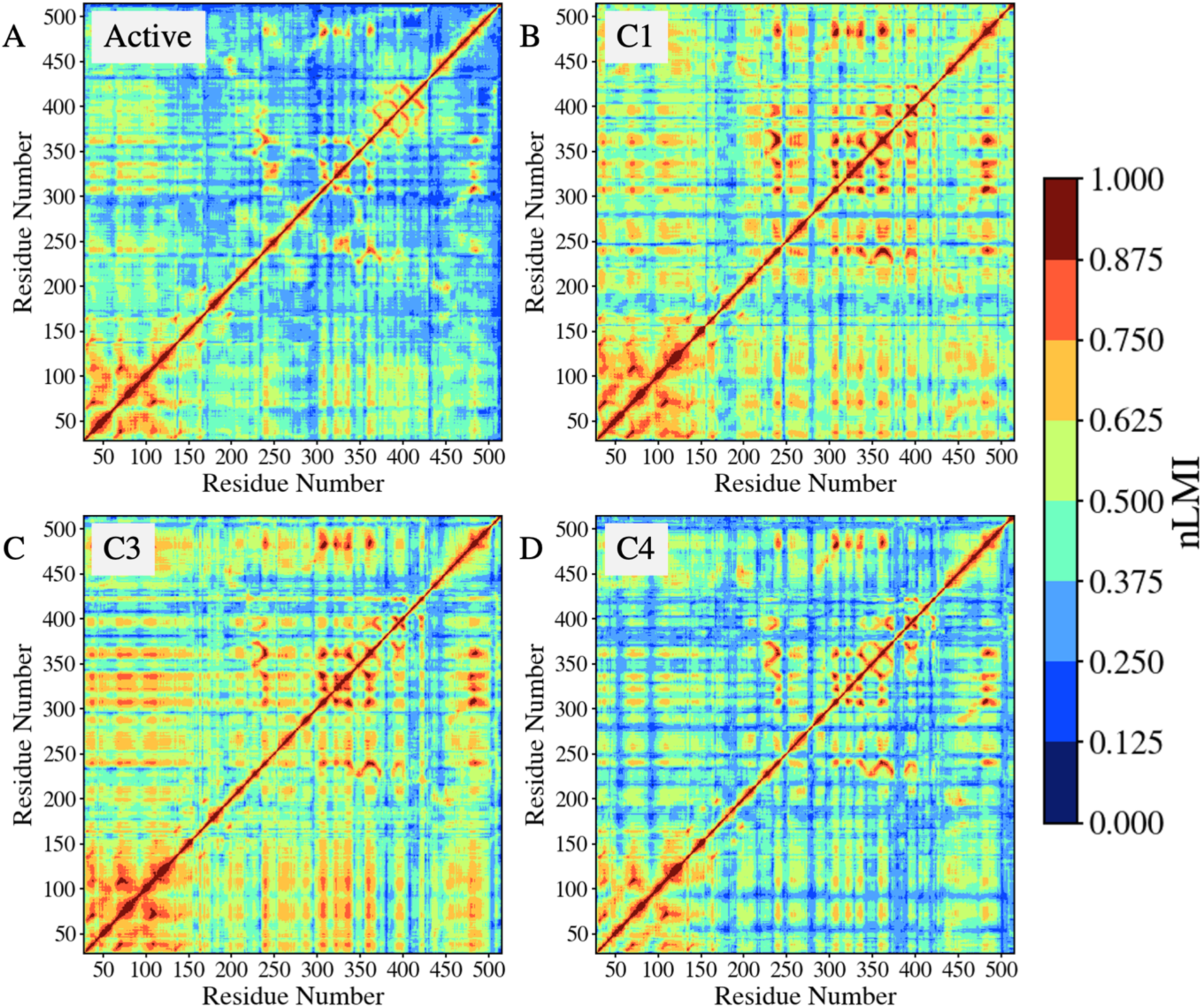
Residue pairwise normalized linear mutual information (nLMI) matrices calculated from MD trajectories using CorrelationPlus package for the active tetrameric state (A) and for G6PDi-1 bound at cavities C1 (B), C3 (C), and C4 (D). nLMI captures correlated motions without the angular dependence that can limit standard dynamical cross-correlation analyses. Here, in the plots, 0 value means that there is no correlation, while 1 means they are completely correlated. Compared with the active state, ligand binding at C1 and C3 produces broader off-diagonal correlation patterns, indicating increased long-range coupling between residues. In contrast, the C4-bound state retains a correlation pattern closer to the active state, with more localized changes.

**Figure S7.**
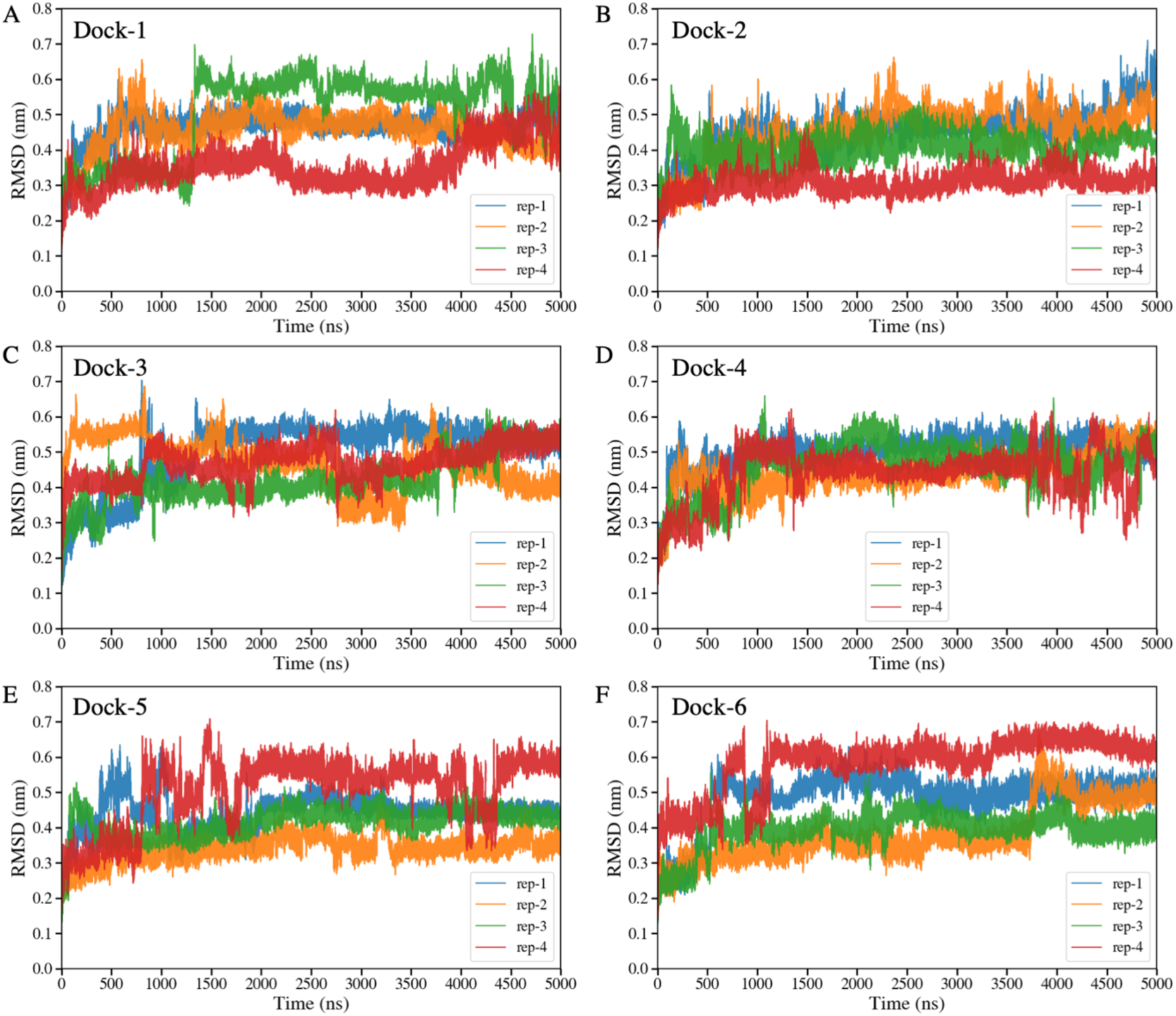
Root-mean-square deviation (RMSD) of the G6PD monomer for the six docked protein-ligand systems (Dock-1 to Dock-6) calculated over the backbone atoms. For each docked pose, four independent replicas of 5μs each were performed.

**Figure S8.**
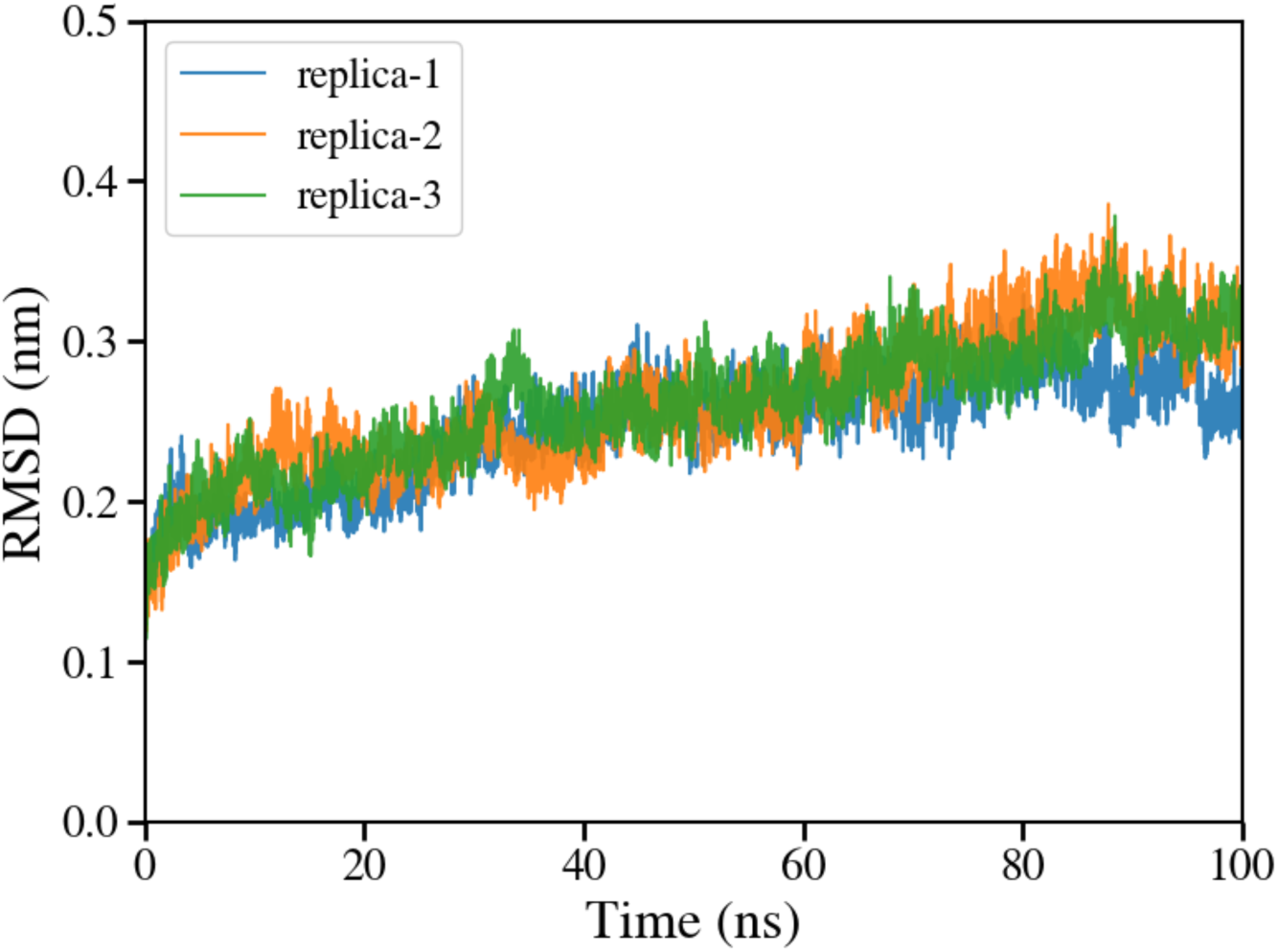
RMSD plot of the G6PD tetramer in active state.

**References**

## Materials and methods

### 1. Experimental details

#### (a) Cell cultures and G6PDi-1 treatment

HepG2 (ATCC-HB-8065) cells were grown in DMEM high-glucose (Euroclone, #ECB7501L, Lot n°EUM01HO) supplemented with 10% fetal bovine serum (FBS) (Euroclone, ECS-5000L, Lot n° EUS53040524), 1% glutamine (Thermo Fisher Scientific, 35050061, Lot n° 3224711), 1% of MEM Non-essential amino acids (SIAL-NEAA-B, Lot n° 23-6365) and 1% penicillin/streptomycin (Euroclone, #ECB3001D, Lot n° EUM01O9). HepG2 cells (2×10^6^ cells/flask) were seeded into 75 cm^2^ flask. The day after cells were treated with 100 µM of G6PDi-1 (HY-W107464, MedChemExpress) for 24 hours, while control cells were treated with the same amount of dimethyl sulfoxide (DMSO).

#### (b) Protein extraction

G6PDi-1 treated and control HepG2 cells were washed with phosphate-buffered saline (PBS) and harvested by trypsinization for 5 min at 37°C. Trypsin activity was neutralized with complete culture medium. Cells were then incubated on ice for 1 hour with Cell Lysis Buffer (1X) (Cell signalling #98039), used to lyse cells under nondenaturing conditions, supplemented with Halt protease and phosphatase inhibitors (Thermo Fisher Scientific, #78444) and 0,9% of Triton X-100 (Sigma-Aldrich #282103). The lysate was centrifuged at 12000 rpm at +4°C for 15 minutes and the supernatant was recovered. Protein concentrations were measured by spectrophotometer.

#### (c) Western blotting analysis

60 ug of cell lysate were loaded in Criterion TGX Stain-Free 10% Precast gel (Biorad, #5671034) and then transferred onto nitrocellulose membrane at 300mA for 90 minutes. The membranes were then incubated in non-fat dry milk 5%, Tween-20 0,05% in TBS at room temperature for 1 hour. The immunoblot was probed with primary and appropriate secondary antibodies against G6PD (Abcam, ab87230). Vinculin was used as endogenous control (Abcam, ab129002). Monomeric and dimeric G6PD bands were detected using a chemiluminescence detection system with the Invitrogen iBright Imaging System (Thermo Fisher Scientific).

### 2. Computational details

#### (a) Binding site prediction and molecular docking protocol

Potential binding site prediction and molecular docking were carried out using the Schrödinger Suite (Schrödinger, 2024) SiteMap module and the molecular modeling package.^1,2^ A monomer structure from the tetrameric crystal structure of human G6PD (PDB ID: 7SNI)^3^ was used as the template. The wild type structure was obtained by back replacing residue Asp200 to Asn. Protein preparation was performed with the Protein Preparation Wizard. Hydrogen atoms were added, hydrogen bonding networks optimized, and the structure subjected to restrained energy minimization. Binding sites on G6PD were predicted and characterized using SiteMap with default parameters. The top six predicted sites (**Figure S2A and Table S1**) were selected for grid generation. Residues within a 5 Å radius of the binding site were set as flexible. Docking calculations were performed using the Glide SP scoring function,^4^ followed by MM-GBSA binding free energy calculations with the OPLS4 force field and the VSGB 2.1 solvation model.^5,6^ The top-ranked docking pose for each cavity was selected based on the docking score and Glide gscore and MM-GBSA binding energy^7^ (**Figure S2B and Table S2**) for subsequent molecular dynamics (MD) simulations.

#### (b) G6PDi-1 molecule parameterization

Parameters of the G6PDi-1 ligand for the MD simulations were generated using the AmberTools24 package in combination with Gaussian16.^8–10^ The initial 3D structure of the ligand was converted into a GAFF2 compatible mol2 file and assigned AM1-BCC charges with Antechamber.^11^ To obtain accurate atomic charges for MD simulations, a three-step quantum mechanical protocol was employed using Gaussian16 package.^8^ First, the ligand geometry was optimized at the Hartree-Fock/PM7 level with tight SCF convergence.^12^ This was followed by a refined geometry optimisation at the B3LYP/def2-SVP level of theory including Grimme’s D3 dispersion correction and an ultrafine integration grid to ensure a stable minimum.^13–15^ Finally, single point electrostatic potential calculations were performed at the HF/6-31G* level using the Merz-Kollman scheme, which provided the input for restrained electrostatic potential (RESP) charge fitting.^16,17^ The RESP derived charges were subsequently assigned to the ligand atoms using Antechamber with the GAFF2 force field.^11^ Any missing bonded or torsional parameters were identified with the parmchk2 utility and compiled into a forcefield modification file. The final outputs, consisting of a GAFF2 mol2 topology and an frcmod parameter file, were then converted into Gromacs format using ParmEd package.^18^

#### (c) Molecular dynamics simulations for G6PDi-1 bound complexes

MD simulations were performed for G6PD in complex with G6PDi-1 docked at six distinct binding sites. For each system (docked complex), four independent replicas were simulated for approximately 5.0 µs each using a 2 fs integration time step. All simulations were performed using GROMACS 2024.2^19^ with the Amber19SB force field^20^ to model the protein, and the TIP3P water model.^21^ Each system was solvated in a periodic box of water molecules and neutralized by the addition of counterions. To mimic physiological conditions, Na⁺ and Cl⁻ ions were added to reach a final salt concentration of 150 mM. Neighbor searching was performed every 20 steps. The PME algorithm was used for electrostatic interactions with a cutoff of 0.9 nm.^22^ A reciprocal grid of 100 x 80 x 96 cells was used with 4th order B-spline interpolation. A single cutoff of 0.93 nm was used for Van der Waals interactions. All bond lengths involving hydrogen atoms were constrained using the LINCS algorithm.^23^ The temperature was maintained at 310 K using the V-rescale thermostat,^24^ and pressure was maintained at 1 atm using the C-rescale barostat,^25^ both under periodic boundary conditions.

All systems were energy minimized using the steepest descent algorithm, followed by a multistep equilibration protocol. For each replica, frames were saved every 10 ps. The total production time generated from these simulations is around 120 µs. All systems were found to be stable during the simulations (**Figure S7**).

#### (d) MD simulations for the wild-type G6PD in tetrameric form

The cryo-EM structure of tetrameric G6PD bound to NADP+ and glucose-6-phosphate (G6P) (PDB ID: 7SNI)^3^ was used to generate the wild type structure. To generate the wild type model, residue 200 was back mutated from Asp to Asn. All simulations were carried out using GROMACS package^19^ with the CHARMM36 force field^26^ and the TIP3P water model.^21^ The parameters for G6P and NADP+ were generated using CGenFF.^27^ Each system was neutralized, followed by energy minimization with the steepest descent algorithm. The system was equilibrated with temperature maintained at 300 K using the V-rescale method^24^ and pressure kept constant at 1 atm using the C-rescale barostat.^25^ Neighbor searching was performed every 10 steps. A reciprocal grid of 96 x 96 x 96 cells was used with 4th order B-spline interpolation. All simulations were performed using periodic boundary conditions, and the particle mesh Ewald method was used for the calculation of electrostatic interactions.^22^ The cut-off distance for electrostatic and van der Waals interactions was set to 1.0 nm. Three independent simulations were run with a time step of 2 fs for 100 ns each, with frames saved at every 10 ps. All trajectories corresponding to the active tetramer state were found to be stable during the simulations (**Figure S8**).

#### (e) MSM estimation and validation

We built a hidden Markov state model (HMSM) to describe how G6PDi-1 transients between multiple cavities on G6PD using PyEMMA.^28,29^ All MD trajectories from the six docked complexes (four replicas each) were used for MSM construction. Seven binding cavities (C1–C7) were defined from contact analysis, and for every frame we computed the minimum distance between the COM of ligand and the COM of residues used to define each cavity. Frames were assigned to a given cavity if the distance was within a cavity-specific cutoff ranging from 0.4–0.6 nm; otherwise, frames were labeled unbound, giving eight observable states in total. The resulting discrete trajectories were passed to PyEMMA to estimate an eight-state HMSM at a lag time of 30 ns. The lag time was selected from implied-timescale analysis (Figure S3A) and the model was further validated by a Chapman–Kolmogorov test. The validated HMSM was then used to derive stationary populations and kinetic observables for ligand exchange between bound cavities and the unbound ensemble (Figure S3B).

